# TRiC/CCT Chaperonin Governs RNA Polymerase II Activity in the Nucleus to Support RNA Homeostasis

**DOI:** 10.1101/2024.09.26.615188

**Authors:** Zlata Gvozdenov, Audrey Yi Tyan Peng, Anusmita Biswas, Zeno Barcutean, Daniel Gestaut, Judith Frydman, Kevin Struhl, Brian C. Freeman

## Abstract

The chaperonin TRiC/CCT is a large hetero-oligomeric ringed-structure that is essential in eukaryotes. While present in the nucleus, TRiC/CCT is typically considered to function in the cytosol where it mediates nascent polypeptide folding and the assembly/disassembly of protein complexes. Here, we investigated the nuclear role of TRiC/CCT. Inactivation of TRiC/CCT resulted in a significant increase in the production of nascent RNA leading to the accumulation of noncoding transcripts. The influence on transcription was not due to cytoplasmic TRiC/CCT-activities or other nuclear proteins as the effect was observed when TRiC/CCT was evicted from the nucleus and restricted to the cytoplasm. Rather, our data support a direct role of TRiC/CCT in regulating RNA polymerase II activity, as the chaperonin modulated nascent RNA production both in vivo and in vitro. Overall, our studies reveal a new avenue by which TRiC/CCT contributes to cell homeostasis by regulating the activity of nuclear RNA polymerase II.

## INTRODUCTION

The molecular chaperone Tailless complex polypeptide 1 Ring Complex (TRiC) or Chaperonin Containing Tailless complex polypeptide 1 (CCT) is an essential eukaryotic chaperonin distantly related to prokaryotic GroEL.^1^ All chaperonins are multi-subunit oligomers forming 2 stacked rings with TRiC consisting of 8 paralogous subunits (Cct1-Cct8) in each ring with an evolutionarily conserved arrangement.^1^ Classically, the chaperonins are considered assembly factors of biological complexes. GroEL was originally identified as an essential host factor for the replication of bacteriophages where it assembles phage capsids (Gro=phage growth; E=growth defect overcome by mutation in phage head protein; L=large subunit).^2–3^ Since these founding studies it has been shown that GroEL also aides in nascent polypeptide folding, prompts the disassembly of protein structures, and is induced to facilitate stress resistance^4–8^. While TRiC also mediates de novo protein folding and complex assembly, it is not stress inducible^9–10^. Overall, the function of eukaryotic chaperonins remains poorly understood.

It is well established that cytoplasmic TRiC promotes the co- and post-translational maturation of ∼10% of new polypeptides including many essential and topologically complex factors including actin, tubulin, many beta-propeller factors such as F-box and G-beta proteins, and cell cycle regulators^6,7,11–14^. Similar to GroEL, a common TRiC/CCT activity is the assembly of protein complexes. For instance, TRiC is needed for the formation of VHL–Elongin BC tumor suppressor structure, the Gβγ dimer, the Bardet– Biedl syndrome protein structure, and the basal transcription factor complex TFIID.^15–18^ However, TRiC does not always contribute to polypeptide folding as is the case when TRiC interacts with gelsolin where the chaperonin regulates the activity of this actin capping/severing factor.^19,20^ Furthermore, TRiC does not fold Orai1 but rather modulates the intracellular trafficking of the native Orai1 membrane protein, which influences calcium signaling kinetics.^21^ Perhaps, it is the multiplex contributions of TRiC to various eukaryotic cellular pathways that make the TRiC chaperonin an essential eukaryotic factor.^22^

Potentially complicating the full understanding of TRiC function, is the division of eukaryotic cells into distinct compartments. While it is clear that prokaryotic-origin mitochondria contain the chaperonin homolog Hsp60, which is more similar to GroEL, whether TRiC functions just in the cytosol or if it has a nuclear role is not understood. TRiC was originally defined as a cytoplasmic machine.^23^ Yet, several studies have reported TRiC subunits in the nucleus and/or associated with nuclear factors including immunofluorescence detection of TRiC subunits in the nucleus.^24–26^ TRiC-P5 was reported in a large complex associated with the nuclear matrix of human lymphocytes and leukocytes, human TCP1 isoforms localize to the nuclear matrix during apoptotic chromatin condensation, and mouse TRiC interacts with heterochromatin during spermatogenesis.^27–30^ High-throughput mass spectrometry studies and selective ribosome profiling analyses have physically connected TRiC to several nuclear factors including components of histone deacetylases (Set3C, Rpd3S, and Rpd3L), histone acetylase (SAGA), chromatin remodelers (SWR-C and SWI/SNF), methyltransferase (Set1C), basal transcription factor (TFIID), RNA metabolism (CCR4-NOT), and RNA polymerase (RNAP II).^6,7,31–35^ While the functional role of TRiC with each of these complexes is yet to be resolved, TRiC has been shown to fold subunits of the SMRT-HDAC3, TFIID, and telomerase complexes and to assemble these subunits into their final structures.^18,36^ Significantly, whether TRiC interacts with these factors in the cytoplasm and/or nucleoplasm isn’t clear. Perhaps notably, the bacterial chaperonin GroEL is associated with RNA metabolism processes including RNA production and mRNA protection.^37–39^ Here, we assessed the functional influence of TRiC on several nuclear pathways and show that TRiC localizes to the nucleus governing RNA polymerase II activity to support nuclear RNA homeostasis.

## RESULTS

### TRiC has a nuclear presence

We began our chaperonin investigation by determining the cellular localization of TRiC in budding yeast cells. We exploited 2 TRiC subunits, Cct2 and Cct6, expressed as GFP fusions within a flexible surface loop of these subunits that do not to interfere with chaperonin assembly and function.^40^ We checked the localization of Cct2-GFP and Cct6-GFP by confocal fluorescent microscopy and biochemical fractionation. Both assays indicated a nuclear presence of TRiC (Figure 1A and 1B). To confirm that these represent the assembled chaperonin, we resolved the nuclear extract preparation by size exclusion chromatography. We observed comparable patterns for Cct2-GFP and Cct6-GFP in nuclear or whole cell extracts with the fully assembled TRiC complex resolving into fraction 2 (Figure 1C). Hence, eukaryotic TRiC is present in the nucleus and is assembled into the 1 MDa holo-chaperonin complex.

**Figure 1.**
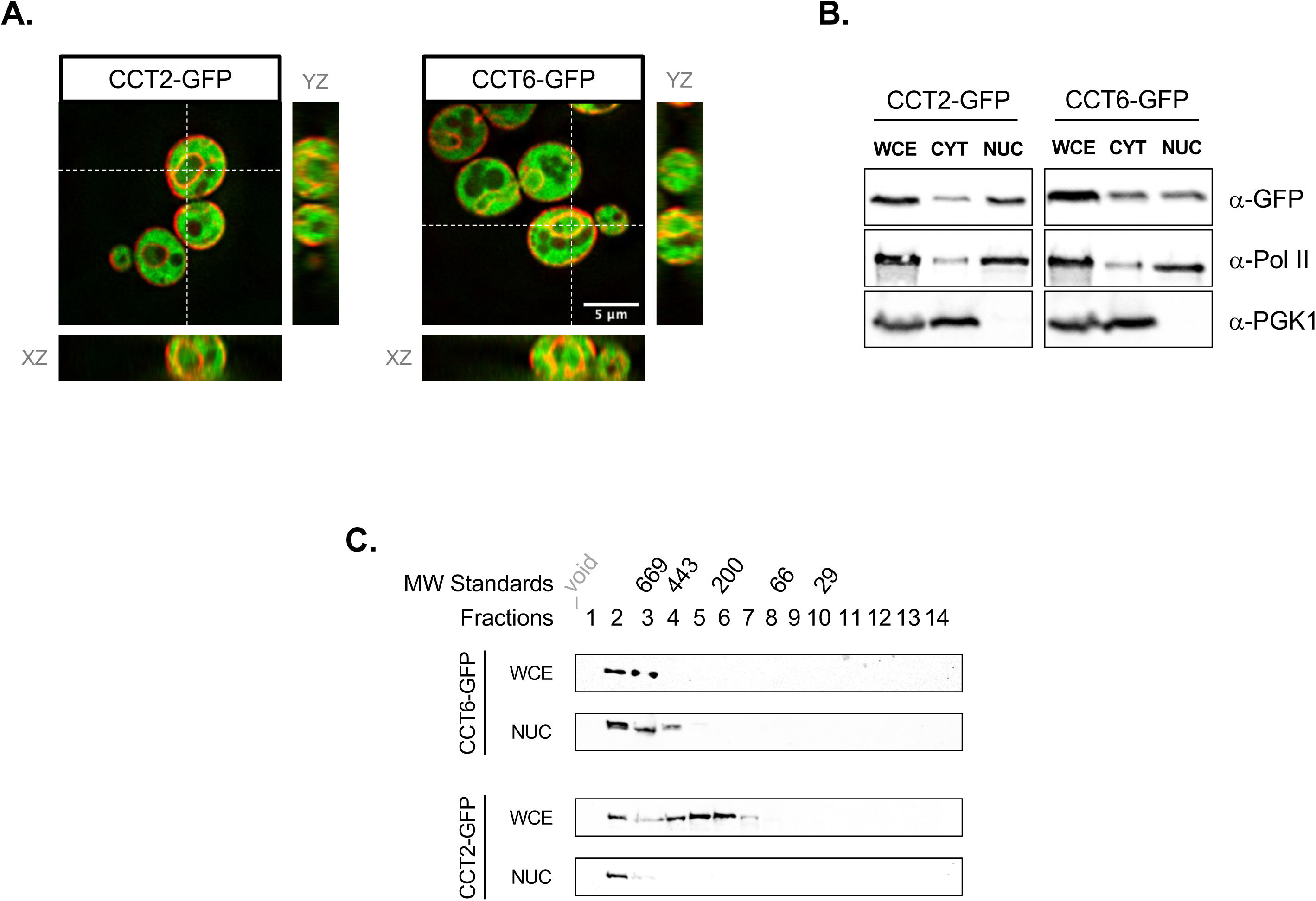
TRiC has a presence in both the cytoplasm and nucleoplasm. (**A**) The cellular localization of TRiC was visualized using yeast expressing GFP-fusions of the Cct2 or Cct6 subunits^40^, as marked. Confocal images were acquired using a DeltaVision OMX. An optical slice (50 μM thick) is presented for each fusion in the x-y planes. Next to each x-y image are the y-z or x-z stacks (500 μM total depth) (**B**) Yeast cells were biochemically separated into cytosolic and nucleosolic fractions. Markers for the nuclear (RNA polymerase α-Pol II) and cytosol (phosphoglycerate kinase 1 α-PGK1) compartments were followed by immunoblot analysis as well as the position of GFP-Cct2 or GFP-Cct6 (α-GFP), as marked. (**C**) To check if GFP-Cct2 and GFP-Cct6 assembled into the chaperonin structure in the nucleoplasm, whole cell and nuclear extracts were resolved by size exclusion chromatography and the position of GFP-Cct2 or GFP-Cct6 were visualized by immunoblot analysis. The relative positions of native protein size standards (units are kDa) are shown above the fractions.

### TRiC inactivation triggers accumulation of divergent noncoding RNA transcripts

To start assessing the influence of the TRiC chaperonin on nuclear events, we examined steady-state RNA transcripts in the presence and absence of active TRiC. To inactivate TRiC we used the temperature sensitive strain *cct1-2* which is fully active at the permissive temperature (30°C) and rapidly inactivates at 37°C.^12,41^ We used an established temperature regime of 4 h at 37°C, which prompts cessation of cell growth that is reversible following a return to a permissive temperature.^41^ In brief, exponentially growing parental (WT) and *cct1-2* yeast were shifted to 37°C for 4 h, RNA was isolated, and RNA-Seq analysis was performed. The most apparent phenotype was the emergence of noncoding/cryptic RNA transcripts including the accumulation of RNAs from bi-directional promoter firing (Figure 2A I1, I2, I3, and I4) and the accumulation of antisense RNAs across open reading frames (Figure 2B and S1A). Of note, there was no apparent correlation between the affected loci and whether these genes were regulated by SAGA or TFIID (data not shown).^42^

**Figure 2.**
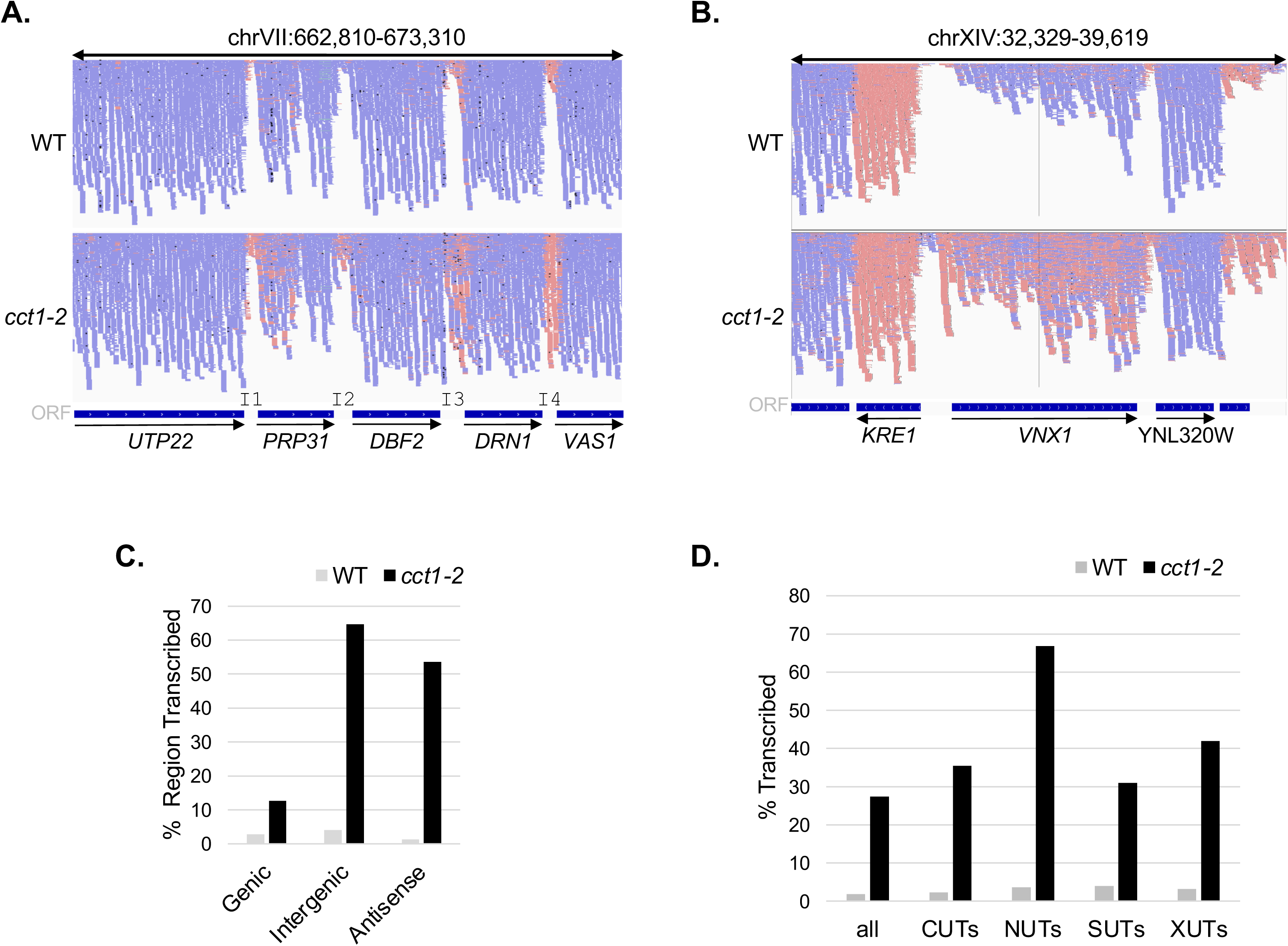
Inactivation of TRiC leads to the accumulation of noncoding RNAs. RNA expression patterns from exponentially dividing wild type (WT) and *cct1-2* yeast exposed to 37°C (non-permissive temperature for *cct1-2*)^41^ for 4 h were determined by deep sequencing (RNA-Seq) and aligning to the yeast genome. Representative patterns illustrating the accumulation of sequence reads in intergenic areas (**A**) or antisense RNAs within an open reading frame (**B**) are shown. The intergenic regions shown in (**A**) were designated intergenic region 1, 2, 3, or 4 (I1, I2, I3, I4). (**C**) TRiC inactivation correlated with a biased increase in the accumulation of cryptic (intergenic and antisense) RNAs. (**D**) The relative representation of aberrant RNAs in WT and *cct1-2* yeast exposed to 37°C for 4 h is shown for known cryptic unstable transcripts (CUTs), Nrd1-unterminated transcripts (NUTs), stable unannotated transcripts (SUTs), and Xrn1-sensitive unstable transcripts (XUTs). *(See also Figure S1)*

**Figure 3.**
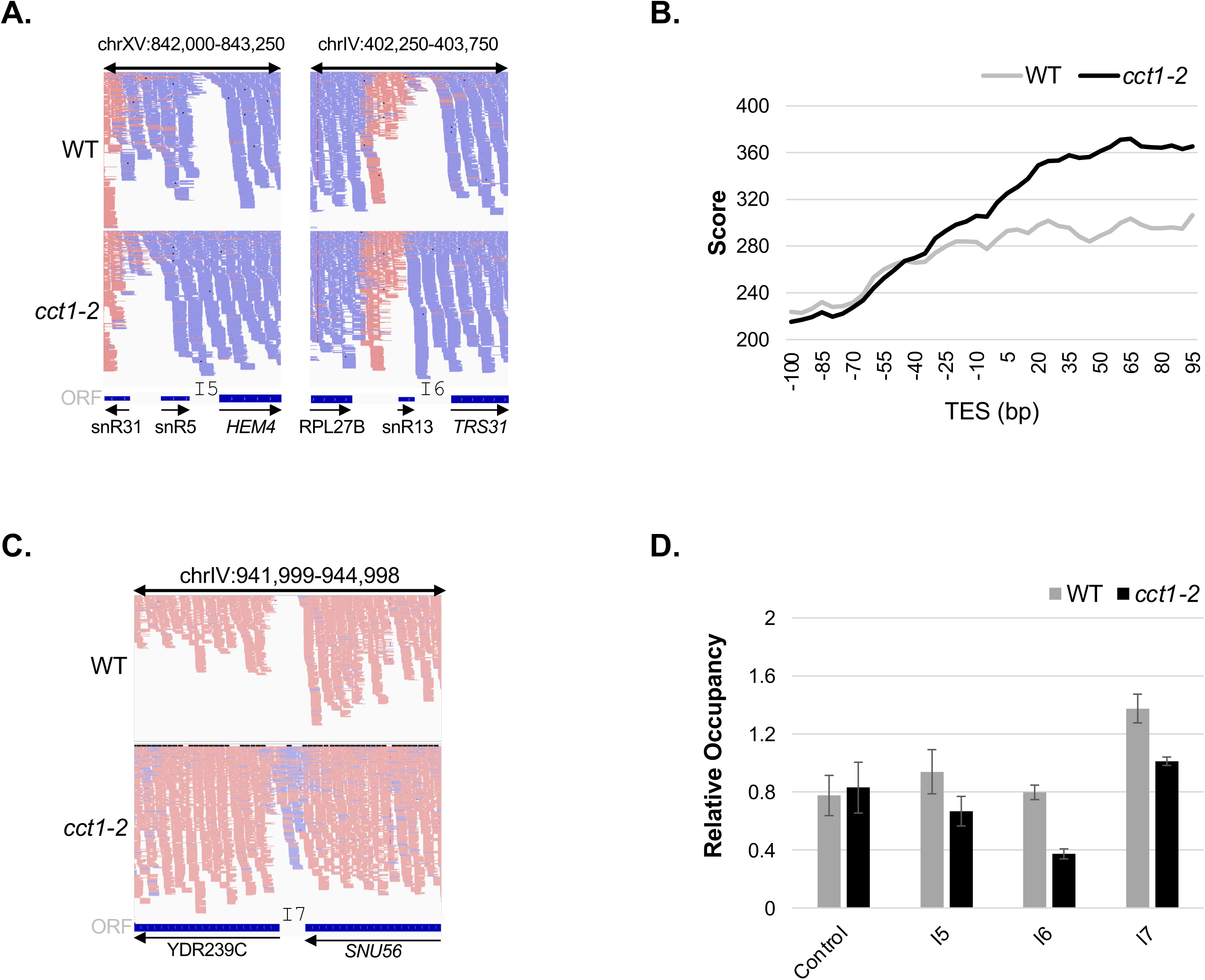
Loss of TRiC prompts read-through transcripts at Nrd1-targeted transcription termination sites. RNA sequence reads from exponentially dividing wild type (WT) and *cct1-2* yeast exposed to 37°C for 4 h were aligned to the yeast genome and the accumulation of the RNAs following the snoRNA encoding genes was checked. The representative snoRNA loci *snR5* and *snR13* are shown (**A**). The intergenic regions downstream of each locus were designated I5 or I6, respectively. (**B**) Meta-analysis of RNA transcript reads around the transcription end sites (TESs) of all snoRNA genes from WT and *cct1-2* yeast exposed to 37°C for 4 h is shown. (**C**) The pattern of RNA accumulation around the TES of the protein encoding gene *SNU56* was assessed in WT and *cct1-2* cells exposed to 37°C for 4 h. (**D**) The relative occupation of the RNA quality control factor Nrd1 was assessed by ChIP at I5, I6, I7, and control (*ACT1*) sites.

**Figure 4.**
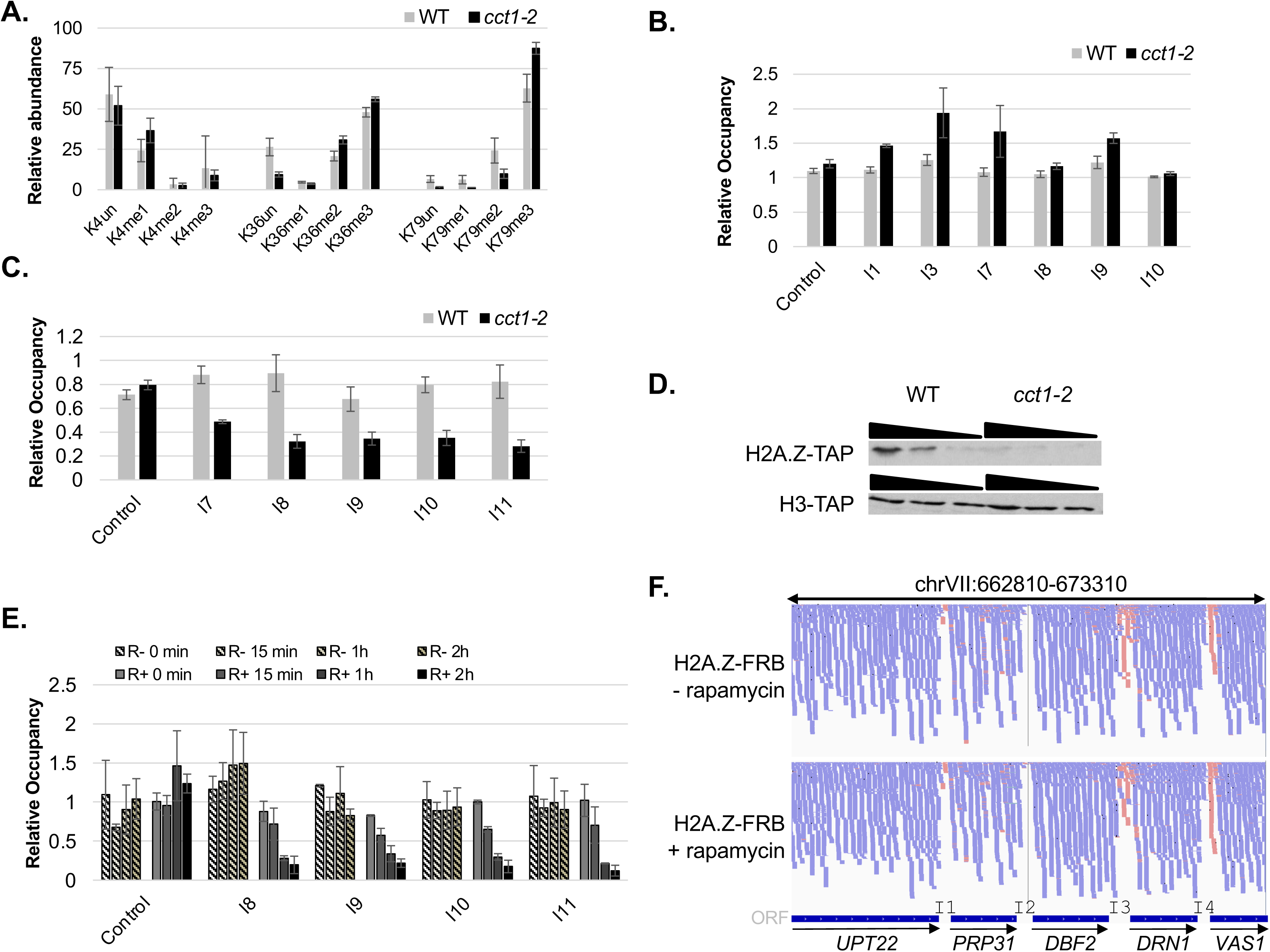
TRiC influence on histone modifications and correlation to cryptic RNAs. (**A**) Mass spectrometry analysis of histone tail post-translation modifications was performed on histone proteins that were isolated from exponentially dividing wild type (WT) and *cct1-2* yeast exposed to 37°C for 4 h. The relative abundance of select H3 modifications are shown. (**B**) The relative levels of H3K79me3 at sites associated with H2A.Z occupancy were determined at control (*ACT1*), I1, I3, I7-I10. (**C**) The relative chromatin occupancy of H2A.Z was determined at control (*ACT1*) and I8-I11 sites in WT and *cct1-2* yeast engineered to express H2A.Z as a TAP-tagged fusion. (**D**) The steady-state levels of H2A.Z-TAP and H3-TAP, as indicated, were checked in WT and *cct1-2* yeast exposed to 37°C for 4 h by immunoblot analysis using an α-TAP antibody and titrations of whole cell extracts (100, 50, 25 μg). (**E**) Using the anchor-away assay, H2A.Z was depleted from the nucleus in engineered WT yeast. The relative chromatin occupancy of H2A.Z at control (*ACT1*), I8-I11 was checked by ChIP following addition of DMSO (striped bars) or Rapamycin (open bars) at the indicated time points. (**F**) The impact of H2A.Z nuclear depletion on RNA patterns was determined using anchor-away and RNA-Seq. Yeast were treated with DMSO or Rapamycin for 2 h, cells were collected, isolated RNA was deep sequenced, and reads were aligned to the yeast genome. *(See also Figure S2)*

While there were changes to steady-state genic RNAs, these were relatively minor with 71 up-regulated (2-GFold; GFold threshold of 0.01)^43^ and 85 down-regulated (50%) genes out of the 6692 open reading frames (ORFs). In general, the activated genes (35/71) were linked to stress response/protein folding and the repressed genes (51/71) were associated with the ribosome/translation. Basically, the changes in sense transcript levels were comparable to mild stress conditions.^44^ Contrasting the limited changes in ORF sense RNAs, loss of TRiC had a large influence on the steady-state levels of cryptic RNAs (i.e., any RNA not representing sense RNA of an ORF) (Figure 2C). To evaluate the accumulated RNAs, normalized RNA-Seq reads along the entire genome were compared from parental and *cct1-2* samples at each nucleotide position. We developed a computational script (Interval Identifier) for the comparison that uses normalized RNA-Seq nucleotide coverages to identify intervals with at least a 4-fold differences in read counts. An advantage of Interval Identifier is that the length of the area with differential coverage between comparison datasets (e.g., parental vs. mutant) depends on the continuity of a 4-fold difference between the adjacent nucleotides. At a threshold where *cct1-2* read-counts were increased ≥4-fold relative to parental (*cct1-2*/WT), ∼9.8% of the genome was associated with noncoding RNAs following TRiC inactivation. In contrast, only ∼0.3% of the genome showed a ≥4-fold increase in parental read-counts compared to *cct1-2* (WT/*cct1-2*). For non-coding transcripts the parental RNA pool covered ∼4% of the upstream intergenic region while *cct1-2* coverage increased to ∼65% of the total intergenic areas. In addition to promoter associated RNAs, 51 ORFs displayed a substantial change in the directionality of transcript production following loss of TRiC (Figure 2B and S1A). Apart from these non-coding RNAs (i.e., bi-directional promoter transcripts and antisense ORF RNAs), there were no clear reference points for where the rest of the cryptic RNAs started and ended since the locations and intensities varied with respect to gene bodies and intergenic regions.

To determine if the diversity of transcripts that are produced in the absence of active TRiC represented known cryptic RNAs, we compared the *cct1-2* pool to previously identified aberrant RNA classes including cryptic unstable transcripts (CUTs), stable unannotated transcripts (SUTs), Xrn1-sensitive unstable transcripts (XUTs), and Nrd1 unterminated transcripts (NUTs). First, we analyzed an established NUT dataset using Interval Identifier and found that it successfully recognized cryptic RNAs (Figure S1B and S1C). Next, we analyzed the WT and *cct1-2* RNA-Seq datasets with Interval Identifier and compared the results to previous reports on CUTs, SUTs, XUTs, and NUTs.^45–48^ The *cct1-2* RNAs overlapped with ∼35% of CUTs, ∼31% SUTs, ∼42% XUTs, and ∼67% NUTs (Figure 2D). Collectively, the known pool of CUTs, SUTs, XUTs, and NUTs covered ∼15% of the *cct1-2* cryptic transcriptome. The changes in these non-coding RNA levels suggest that a TRiC-dependent breakdown in the RNA quality control process accounts for much of the cryptic transcripts present in *cct1-2* cells.

### TRiC inactivation leads to Nrd1-dependent termination defects at snoRNA loci

To check whether TRiC is involved with a known RNA quality control component, we examined sites associated with the RNA quality control factor Nrd1. Studies on Nrd1 have shown that it is involved in many classes of aberrant nuclear RNAs and is therefore a key constituent for the global surveillance of the transcriptome.^48,49^ Since the binding motif for Nrd1 is enriched in snoRNAs^48^, we first examined these loci. In general, snoRNA encoding genes displayed increased readthrough beyond the transcription termination sites (Figure 3A I5 and I6). For almost half of the snoRNA genes (30 out of 68) the level of readthrough transcripts was ∼1.5-fold higher in *cct1-2*, yet the level of RNA prior the termination site was comparable (Figure 3B). Readthrough transcripts were also apparent at the non-snoRNA gene, *SNU56*, which is also targeted by Nrd1 (Figure 3C). Of note, divergent RNAs from read through of *SNU56* and from bidirectional promoter firing of YDR239C are both present in intergenic region I7 (Figure 3C). All of these loci displayed a mild decline in Nrd1 occupancy at the 3’ ends as measured using the chromatin immunoprecipitation (ChIP) assay (Figure 3D). Yet, given the difference in the magnitude of the cryptic RNA phenotype versus the mild change in Nrd1 occupancy, we concluded Nrd1 impairment does not explain the observed RNA changes occurring after TRiC inactivation. We therefore explored the influence of TRiC on other nuclear pathways including chromatin-associated features.

**Figure 5.**
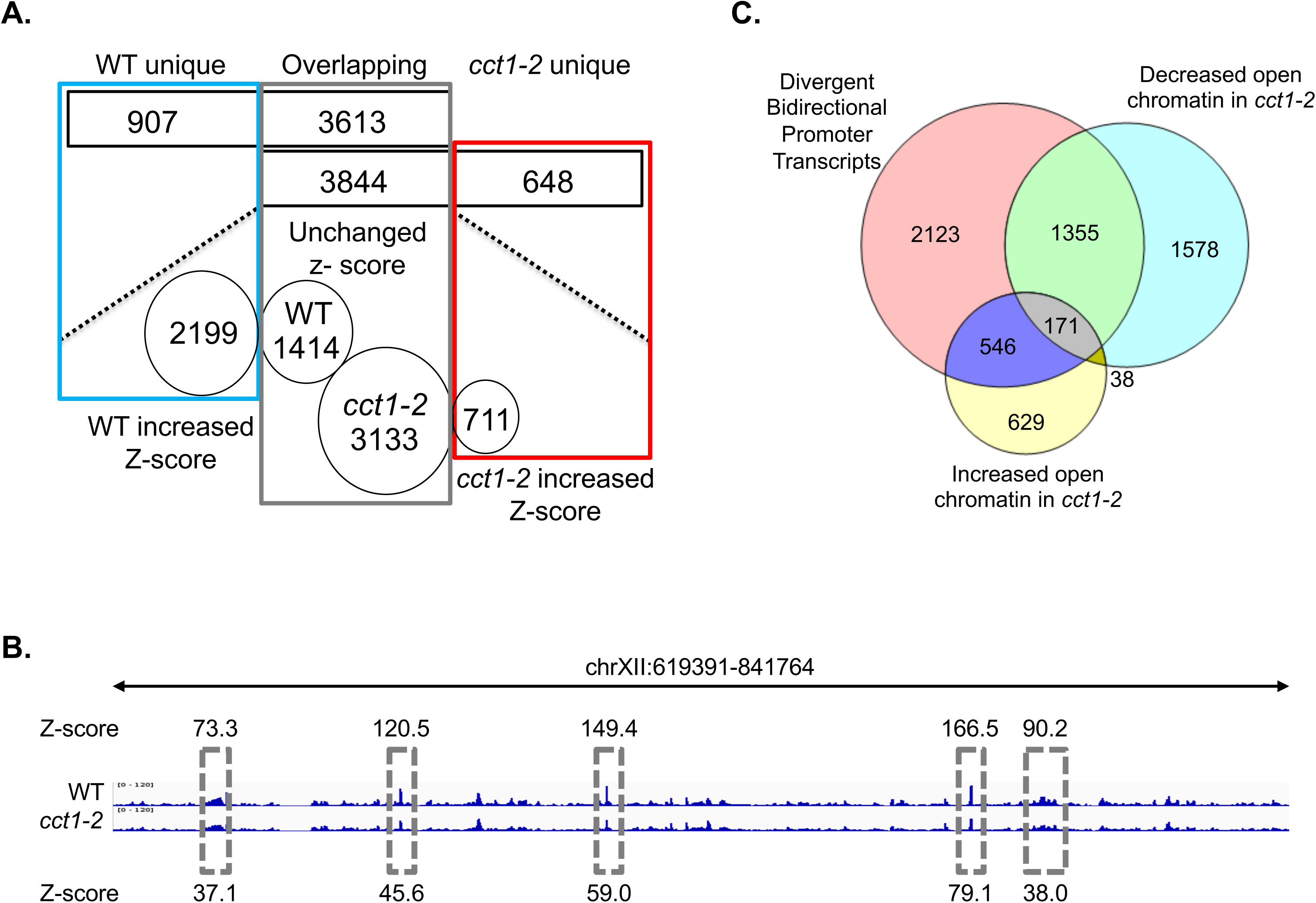
Local chromatin structure is mildly altered following loss of TRiC. (**A**) Chromatin accessibility was probed by DNase I cleavage in exponentially dividing wild type (WT) and *cct1-2* yeast exposed to 37°C for 4 h. The blue box highlights the number of DNase I hypersensitivity sites (DHSs) unique to WT, the grey box is the overlapping DHSs, and the red box marks the DHSs unique to *cct1-2* after 4 h at 37°C. The circled numbers indicate the DHSs with either unchanged or increased Z-scores (hypersensitivity/openness) in either WT or *cct1-2*, as marked. (**B**) A representative section of the genome is shown with the hatched grey boxes denoting DHSs that decreased in *cct1-2* yeast. The DHS Z-scores are provided for each TRiC-dependent DHSs. (**C**) Venn diagram illustrating the intersections between cryptic transcripts associated with bidirectional promoter firing and either decreased or increased open chromatin following TRiC inactivation.

**Figure 6.**
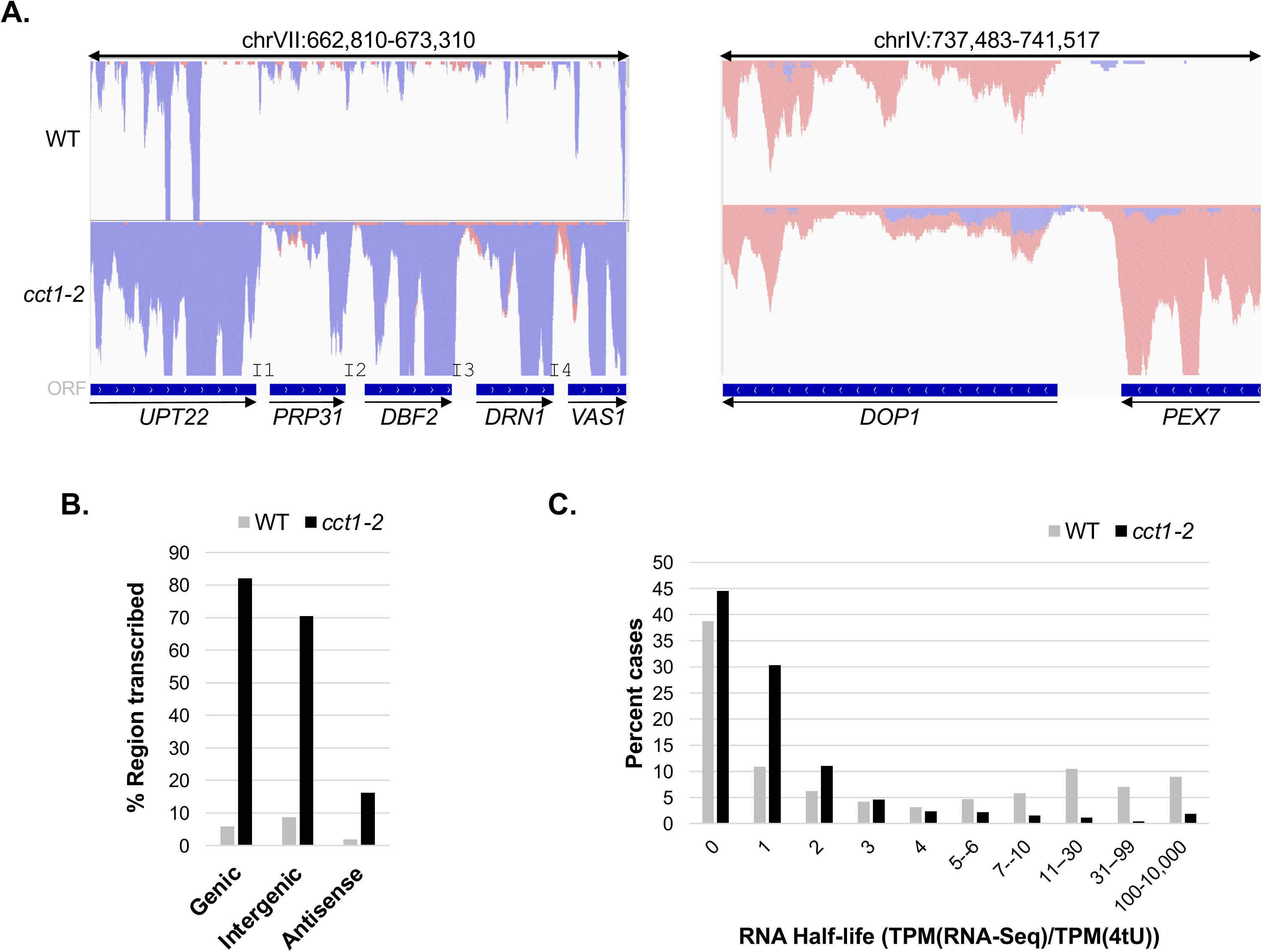
Production of nascent RNAs is TRiC-dependent. (**A**) The relative levels of active transcription in exponentially dividing wild type (WT) and *cct1-2* yeast exposed to 37°C for 4 was monitored by 4tU-Seq. For illustrative purposes, 2 different areas of the genome are shown with the sequences reads aligned to the yeast genome from samples prepared from WT or *cct1-2* yeast exposed to 37°C for 4 h. (**B**) TRiC inactivation increased the production of nascent RNA transcripts across the genome regardless of RNA type (i.e., genic, intergenic, or antisense). (**C**) Relative RNA half-lives in WT and *cct1-2* cells exposed to 37°C for 4 h were calculated as TPM(RNA-Seq)/TPM(4tU-Seq) where TPM=transcripts per million.

### Influence of TRiC on epigenomic marks

The TRiC chaperonin has been linked to numerous chromatin modifiers including SAGA, NuA4, RPD3S/L, SET3C, SET1C, HDA1, HST3/4, and SPT10.^31–35,50^ TRiC is minimally needed for the proper assembly of the SMRT-HDAC3 and TFIID complexes.^18,36^ We thus examined whether TRiC has an active role in modulating the epigenomic landscape that could account for the large increase in steady-state cryptic RNAs. Using standard procedures, we isolated histones from exponentially growing parental (WT) and *cct1-2* cells incubated for 4 h at 37°C and interrogated 23 different histone posttranslational modifications (PTMs) by mass spectrometry.^51^

Perhaps unexpectedly, the only observed significant change to the histone code was an ∼25% (±8% SD) increase of H3K79me3 in *cct1-2* cells (Figure 4A and S2A-C). The H3K79me3 PTM often demarcates nucleosomes at or near actively transcribed loci and H3K79me3 has been implicated in pervasive extragenic transcription.^52–54^ Of note, ∼29% of the sites displaying increased steady-state transcripts in *cct1-2* overlapped with established H3K79me3 peaks (data not shown).^54^ We did find a small increase in H3K79me3 at sites subject to elevated cryptic transcripts (Figure 4B). However, not all sites with cryptic RNAs correlated with elevated H3K79me3 levels (Figure 4B I8 and I10). The disparity in the magnitude of the TRiC-dependent differences between epigenomic changes and the increased levels of cryptic RNAs upon TRiC inactivation suggests other/additional factors contribute to the accumulation of the non-coding RNAs.

As H3K79me3 has been reported to be mutually exclusive with H2A.Z histone variant^55^, we next examined the correlation between H2A.Z and pervasive RNA production. We found that H2A.Z occupancy declined ∼2-fold at sites having elevated divergent RNAs in *cct1-2* (Figure 4C). The reduction in H2A.Z occurred in a time dependent manner after TRiC inactivation (Figure S2D). Of note, the TRiC-dependent decrease in H2A.Z chromatin association appears to be independent of H3K79me3 since a similar change in H2A.Z occupancy was observed in *dot1*Δ yeast following TRiC loss (Figure S2E)—Dot1 is the lone H3K79 tri-methyltransferase in yeast.^57^

We found that the steady-state levels of the H2A.Z protein declined in *cct1-2* at the restrictive temperature with no apparent change in H3 protein (Figure 4D), which suggests the H2A.Z protein relies on the TRiC chaperonin for stability. Of note, deletion of the H2A.Z encoding gene *HTZ1* further sensitized *cct1-2* yeast to increased temperature thereby demonstrating a clear physiological link between these factors (Figure S2F). Given the connections between TRiC and H2A.Z, we tested if a reduction of H2A.Z at gene promoters instigated an accumulation of cryptic RNAs comparable to TRiC inactivation. To trigger the loss of H2A.Z in chromatin, we engineered an H2A.Z anchor-away yeast strain. In brief, anchor-away allows the controlled removal of a protein from the nucleus by taking advantage of the Rapamycin-dependent interaction between FKBP12 and the FKBP Rapamycin-Binding domain (FRB) of TOR1 in addition to the flow of ribosomal proteins from the nucleus to the cytoplasm.^58^ Following rapamycin addition, we observed a time-dependent depletion of H2A.Z from chromatin as measured by ChIP (Figure 4E). Despite the H2A.Z reduction, there was no significant change in the accumulation of divergent RNAs, as determined by RNA-Seq (Figure 4F). Overall, these experiments indicate that H2A.Z is likely a client of TRiC, yet the TRiC-dependent loss of H2A.Z does not account for the increase in steady-state levels of cryptic transcripts upon TRiC inactivation.

### Impact of TRiC inactivation on chromatin accessibility

Our prior work on the roles of molecular chaperones in the nucleus showed that both Hsp90 and p23 impact chromatin accessibility.^59–61^ Hence, we evaluated whether the TRiC chaperone might have a similar influence by mapping the global DNaseI hypersensitivity sites (DNaseI-Seq).^59,62^ We observed that TRiC does not modulate the total quantity of open chromatin regions across the genome. The number of DNaseI hypersensitive sites (DHSs) was comparable following TRiC inactivation (parental DHSs=4520 and *cct1-2* DHSs=4492). Yet, there were changes to the locations of the DHSs (∼20% of parental DHSs are lost at the restrictive temperature with an ∼16% gain in DHSs specific to the *cct1-2* background) (Figure 5A). Also, the average DHS Z-score, which is a measure of DNase I cleavage level^63^, reduced from 19.5 in parental cells to 13.2 in *cct1-2* (32% reduction) indicating a mild reduction in chromatin accessibility following loss of TRiC (Figure 5B). These experiments show that TRiC can influence chromatin accessibility since access was reduced mildly. Yet, the changes in DHSs (i.e., decreased size, increased, or unique to background) did not correlate with the production of cryptic RNAs (Figure 5C). For example, ∼50% of the divergent RNAs associated with bidirectional promoter firing did not display a change in accessibility. To this point, we conclude that the observed increases in cryptic RNAs following TRiC inactivation are not accounted for mechanistically by changes in RNA quality control, histone modifications, or chromatin accessibility (Figures 2-5).

### Loss of TRiC triggers a significant increase in nascent RNA production

We next focused on the possible role of TRiC in the process of RNA biogenesis. The steady-state levels of any RNA transcript is set by multiple events including RNA synthesis and decay pathways.^64^ To evaluate the global dynamics of nascent transcript production we exploited metabolic labeling of RNA with the nucleotide analog 4-thioracile (4tU) followed by high-throughput sequencing (4tU-Seq).^65,66^ In brief, 4tU-Seq allows the relative contributions of synthesis and decay rates to steady-state RNA levels to be appraised.^66^ Hence, we evaluated nascent RNAs in parental and *cct1-2* cells using 4tU-Seq.

Following a 4 h incubation at 37°C, parental and *cct1-2* cells were exposed to 4tU for 5 min to label the nascent transcripts. RNA isolated from labeled and unlabeled (control) cells were conjugated to biotin with a thiol-reactive reagent, the biotinylated RNA was affinity purified with streptavidin, and the isolated RNA was sequenced. Significantly, we observed a dramatic increase in nascent RNA production in *cct1-2* yeast (Figure 6A). At a threshold of ≥4-fold for *cct1-2*/WT, ∼22.8% of the genome displayed higher levels of nascent RNAs whereas only ∼0.4% of the genome had levels ≥4-fold when comparing WT to *cct1-2*. Relative to the steady-state RNAs (Figure 2), nascent transcripts cover more of the genome (∼9.8 vs. 22.8%) and are ∼7.3-fold more abundant. In contrast to the biased changes in intergenic and antisense transcripts within the steady-state RNA pool of *cct1-2* (Figure 1), the nascent RNAs were elevated across the genome in *cct1-2* (Figure 6B). As might be anticipated, given the broad genomic expression pattern of the *cct1-2* transcripts, the *cct1-2* nascent RNAs overlapped well with all classes of known cryptic RNAs including CUTs, SUTs, XUTs, and NUTs (data not shown). Thus, the increased production of nascent RNAs is contributing to the accumulation of non-coding RNAs.

In addition to evaluating nascent RNA production, the 4tU-Seq data in conjunction with RNA-Seq can be used to gauge RNA degradation rates.^65,67^ In brief, RNA half-lives are proportional to the normalized number of RNA molecules (RNA-Seq) divided by the level of normalized, actively produced RNAs (4tU-Seq). Analysis of the parental (WT) and *cct1-2* datasets revealed that the WT RNA pool has longer-lived transcripts (i.e., parental pool has higher levels of RNA species with a half-life estimator value of ≥4) and shorter-lived transcripts are preferentially found in *cct1-2* (i.e., *cct1-2* pool is enriched for RNAs with half-life estimator values ≤4) (Figure 6C). Hence, the loss of TRiC resulted in an increased production of nascent RNAs but also shortened RNA lifespan. Likely, the increased degradation rate is the cell’s attempt to restore RNA homeostasis by countering the increased production rate. We suspect that the increase of cryptic RNAs in the steady-state pool (Figure 2), results from a preferential activity of the RNA quality control machinery on coding RNAs, which would reduce the stable output of coding RNAs to near normal but prolong the lifetime of non-coding RNAs. This could be due to either RNA quality control being overwhelmed by the increased level of substrate or by a higher affinity towards coding RNAs. Regardless of the mechanism, these experiments indicate a major outcome of TRiC loss is the increase in the global production of RNAP II-dependent nascent RNAs.

### Nuclear depletion of TRiC triggers increased noncoding RNA accumulation

We next asked if the effect of TRiC loss on RNA transcript production involves its cytoplasmic function or whether it is due to nuclear role of TRiC. Initially, we attempted to use the anchor-away approach to remove TRiC from the nucleus but, unfortunately, our attempts to use various TRiC subunits as FRB fusions were unsuccessful. Instead, we exploited our existing Cct2-GFP and Cct6-GFP strains and transformed these cells with a *GAL1*-driven expression vector for a GFP nanobody that also has 3 nuclear export signals (NES-GFP-Nanobody). Following the addition of galactose to the media, we found a time-dependent nuclear depletion of Cct2-GFP or Cct6-GFP (Figure 7A).

**Figure 7.**
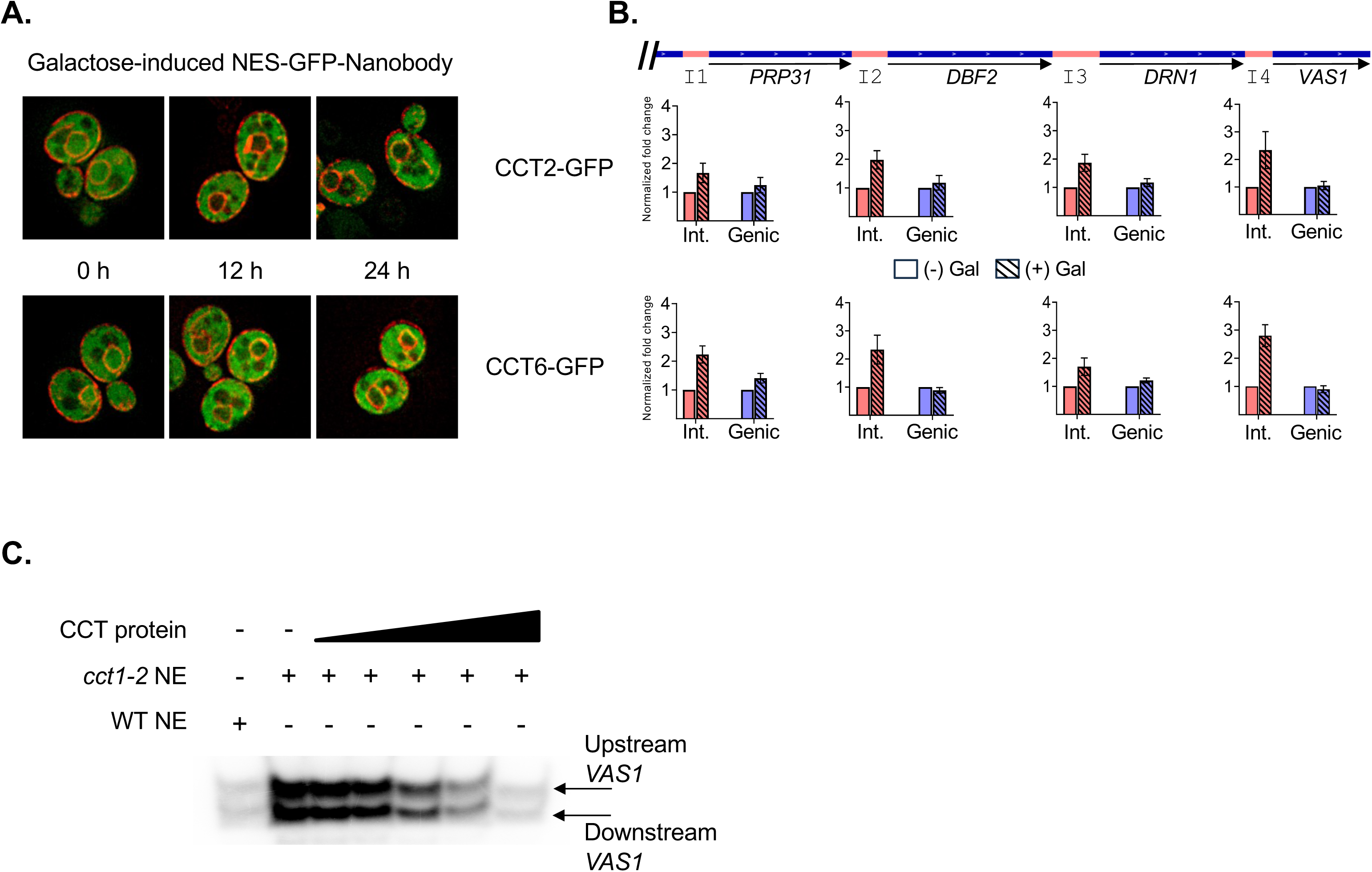
TRiC directly regulates RNA polymerase activity. (**A**) The cellular localization of TRiC was visualized using yeast expressing the Cct2 or Cct6 subunit as a GFP-fusions and a mCherry-Pho88 fusion to mark the outer and nuclear membranes.^40,83^ The yeast transformants also carried an expression vector for a GFP-Nanobody with nuclear export signals (3x NES) under the control of the *GAL1* promoter. To trigger nuclear depletion of TRiC, yeast growing exponentially in selective simple raffinose-based media were supplemented with galactose. The GFP signal was visualized at 0, 12, and 24 h post galactose addition. (**B**) Yeast expressing GFP-fusions of either Cct2 or Cct6, as marked, were transformed with the NES-GFP-Nanobody. Exponentially growing cells were supplemented with galactose, incubated for 24 h, and RNA was isolated. The production of intergenic (Int.) or genic (Genic) RNA was monitored by RT-PCR using select primers. The normalized intergenic signal is represented by pink bars and normalized genic signal by blue bars. (**C**) In vitro transcription reactions were performed using nuclear extract (NE) prepared from exponentially dividing wild type (WT) or *cct1-2* yeast exposed to 37°C for 4 h, as marked. The *cct1-2* NE was supplemented with purified, recombinant TRiC (5, 10, 25, 50, 100 nM final), as indicated. The production of genic (downstream *VAS1*) or intergenic (upstream *VAS1*) RNA was detected by primer extension using select oligonucleotides.

Significantly, the nuclear depletion of TRiC led to an increase in steady-state intergenic RNAs but not genic species (Figure 7B). Hence, the presence of TRiC in the nucleus is required to regulate the production of RNAs and long-term nuclear depletion leads to an imbalance in the intergenic vs. genic RNAs.

### TRiC regulates RNA polymerase II activity in vitro

To assess whether TRiC acts directly on RNAP II activity we employed an in vitro transcription assay. We tested the in vitro production of RNA from the *VAS1* promoter, which displayed increased noncoding RNA levels associated with bidirectional promoter firing in vivo (I4 in Figure 2A and 6A). Parental and *cct1-2* yeast were incubated 4 h at 37°C prior to preparing nuclear extracts. A biotinylated *VAS1* promoter DNA template was immobilized on magnetic beads and the bidirectional production of RNAs from the *VAS1* transcription start site (TSS) was visualized by standard primer extension assays with oligonucleotides select for upstream or downstream RNAs. The *VAS1* promoter fired pervasively producing transcripts in an orientation independent manner with the *cct1-2* nuclear extract creating ∼10-fold more RNA in both directions (Figure 7C). Importantly, titration of purified, recombinant TRiC into the *cct1-2* nuclear extract was sufficient to reduce the activity of RNAP II, in a dose dependent manner, to the level of RNAP II in WT nuclear extract (Figure 7C). These results indicate that TRiC can directly control the activity of RNAP II to achieve a WT transcriptional output that balances RNA homeostasis in the nucleus.

## DISCUSSION

A central feature of proteostasis is the recognition and manipulation of protein conformations by molecular chaperones to support and regulate client form and function.^68^ Typically, the emphasis is on nascent polypeptide biogenesis and damaged protein triage. However, chaperones can also regulate the activity of mature cellular proteins thereby conferring a mechanism for recurrent protein regulation.^69^ Here, we show that TRiC has a function in the nucleus that is independent of its cytoplasmic role. Nuclear TRiC is required to modulate RNA polymerase II activity to maintain a proper level of transcriptional output and balance RNA homeostasis. As RNAP II is reiteratively used to produce RNA transcripts, cycling this enzyme complex through a central molecular chaperone (i.e., TRiC chaperonin) would link the proteostasis process to the production of RNA transcripts. Essentially TRiC would be serving as an RNAP II rheostat conferring a persistent regulatory influence based on chaperonin availability. Under conditions where TRiC activity is occupied with non-RNAP II clients, the production of RNAP II transcripts would increase. For instance, during physiological conditions demanding a higher synthesis rate (e.g., organism development or cell differentiation), TRiC might be heavily engaged in the biogenesis of nascent polypeptides, assembly of protein complexes, and/or obligate folding of the abundant cytoskeletal proteins actin and tubulin.^70^ Occupying TRiC with other duties would foster higher production of RNAP II transcripts that, in turn, would allow increased protein synthesis to support the cell diversification. Cell demand would selectively stabilize requisite protein coding transcripts. In our model, TRiC is a key determinant in balancing proteostasis and mRNA production.

Classically, the modulation of proteins by a chaperonin is thought to occur within the inner cavity of the 2 stacked rings forming the TRiC structure.^1^ As the size limitation of this folding chamber is ∼70 kDa, this raises an obvious concern on how TRiC might regulate the >500 kDa RNAP II complex. Yet, TRiC has been shown to facilitate the folding of proteins larger than 70 kDa including the ∼110 kDa spliceosome subunit Snu114 where TRiC partially encapsulates the carboxyl-domain of Snu114 to prompt its folding while the amino-terminal domain is independent of TRiC.^71^ Hence, TRiC may regulate RNAP II by engaging a protruding domain of a subunit of the assembled RNA polymerase enzyme complex.

Significantly, TRiC/CCT dysregulation has been linked to various ailments including numerous cancers, neurodegenerative diseases, myocardial infarction, and Down’s syndrome.^72^ It had been assumed that any malady manifesting from a TRiC defect would result in a protein health deficiency, yet our work opens the possibility that TRiC shortcomings might trigger transcriptome-related problems. Of note, recent advances have implicated noncoding RNA transcripts as cancer determinants.^73^ For example, Downstream-of-Gene (DoG) noncoding transcripts, which are a class of long noncoding RNAs (lncRNAs), contribute to carcinogenesis and correlate with patient survival.^74,75^ Through undefined mechanisms, DoG RNA levels increase in response to physiological stress including heat shock, osmotic stress, and viral infection.^76–79^ As all of these conditions can impinge upon TRiC activity^72^, it is intriguing to consider whether the stress-related expression of DoG RNAs is TRiC-dependent.

With the onset of genomic research, a myriad of noncoding transcripts, including promoter-associated pervasive RNAs and natural antisense transcripts (NATs), were found in organisms from bacteria to humans.^80,81^ In humans, ∼55% of all promoters fire bidirectionally with little bias in orientation and ∼40% of all open reading frames are transcribed in both directions within a cell population.^80^ Despite the commonality of divergent RNAs, finding a connection to the TRiC chaperonin was unexpected. Nonetheless, our open investigation into the nuclear roles of TRiC converged on its influence on RNAP II activity (Figures 2, 6, and 7). TRiC inactivation led to the accumulation of different steady-state noncoding RNAs including upstream transcripts consistent with pervasive bidirectional promoter firing, antisense RNAs within open reading frames, and read through transcripts (Figure 2 and 3). A major contributing factor to the production of these noncoding RNAs is likely the increased RNAP II activity found in the absence of TRiC both in vivo and in vitro (Figure 6 and 7). Interestingly, the TRiC-dependent enhanced production of RNA from bidirectional promoter firing showed little orientation bias in vitro (Figure 7C). Yet in vivo only the noncoding RNA accumulated (Figure 2 and 7B), suggesting a mechanism to control the rise in protein coding RNAs. Minimally, the act of translating an mRNA is sufficient to destabilize a transcript^82^ or, perhaps, an additional pathway exists to selectively reduce the levels of coding RNAs to normal. No matter the destabilizing event, we suspect that the increased load of coding RNA is sufficient to saturate the RNA quality control machinery thereby increasing the lifespan of the observed noncoding RNAs. Overall, our work identifies an important new role for the TRiC chaperonin in the control of RNA polymerase II activity in the nucleus to maintain RNA homeostasis.

## STAR Methods

### Resource Availability

#### Data and code availability

- All DNA sequencing, mass spectrometry (PRIDE), and imaging data (Mendeley Data) generated in this study have been deposited and are publicly available as of the date of publication. Accession numbers for the RNA-Seq, DNaseI-Seq, 4tU-Seq are GSE274453, GSE274454, GSE274455.
- The Interval Identifier code has been deposited at Zenodo (https://zenodo.org/records/7804010) and is publicly available.
- Any additional information required to reanalyze the data reported in this paper is available from the lead contact upon request.

### Experimental Model and Study Participant Details

*Saccharomyces cerevisiae* cultures were grown in complete (YPD) or minimal media supplemented with the required amino acids at 30°C unless otherwise noted. The main yeast strains (WT and *cct1-2*) were previously described.^41^ TAP-tagged variants were produced by standard recombinant methods by introducing the fusion tag into either parental (WT) or *cct1-2* yeast.^84^ The yeast expressing the CCT subunits Cct2 and Cct6 as GFP fusions were previously described.^40^

### Method Details

#### Cell fractionation for size exclusion chromatography

Cell fractionation was performed as described^85^ with the following modifications. Briefly, cells were treated with 0.1% sodium azide to halt nuclear transport before spheroplasting. An aliquot was removed before lysis by Dounce homogenization representing the whole cell extract. Upon centrifugation, 5% of total cell amount was taken from the supernatant (Cytoplasmic fraction) which was concentrated using TCA (10%) precipitation. The pellet was resuspended in 8% PVP with 0.025% Triton X-100 in 20mM potassium phosphate buffer pH 6.5, 1mM MgCl2, 1mM DTT and protease inhibitors and incubated on ice for 10 min. Nuclei were obtained by centrifugation at 10000xg for 5 min. Cell fractionation was visualized via immunoblotting using rabbit anti-GFP (sc-8334), mouse anti-PGK1 (Invitrogen ab113687) and mouse anti-PolII(CTD) (Santa Cruz sc-47701). Whole cell extract and nuclear fractions also were resolved by size exclusion chromatography using a Superdex-200 column (Cytiva). After passage of the void volume, 1 mL fractions were collected, precipitated with TCA (10% final), and the samples were analyzed by immunoblot using a rabbit anti-GFP (sc-8334).

#### RNA-Seq

Total RNA was isolated from WT or *cct1-2* yeast using Yeast RiboPure RNA purification kit (Ambion) from cells grown to log phase at 30°C and then shifted to 37°C for 4 h. The libraries were prepared using TruSeq Stranded mRNA kit (Illumina) according to manufactures instructions, the samples were sequenced on an Illumina HiSeq2000, and processed with Casava 1.8.2. The trimmed reads were aligned to the sacCer3 reference genome (UCSC) with STAR and differences in gene expression were determined with GFold.^43,86^

#### DNase-Seq

The DNase-Seq protocol was adapted from an established method^56^, as described previously.^59^ In brief, nuclei were prepared from WT or *cct1-2* yeast by Dounce homogenization of spheroplast cells from cells grown to log phase at 30°C and shifted to 37°C for 4 h. Nuclei were treated with DNaseI at room temperature, DNA was isolated, resolved on a 1% agarose gel, the DNAs in the size range of 100-650 bp were excised, DNA was isolated using a QIAprep Spin Miniprep columns (Qiagen), and sequenced on an Illumina HiSeq2000. DNaseI hypersensitive sites were identified with hotspot and DNA footprints as described.^59,61,62,87,88^

#### Histone tail post-translation modification mass spectrometry

Total histones were isolated from WT or *cct1-2* yeast grown to log phase at 30°C and shifted to 37°C for 4 h using an established acid histone extraction protocol.^89^ Nuclei were prepared from the collected cells using Nuclei Isolation Buffer (15 mM Tris (pH 7.5), 60 mM KCl, 15 mM NaCl, 5 mM MgCl_2_, 1 mM CaCl_2_, 250 mM Sucrose, 1 mM DTT, 2 mM sodium vanadate, 1x protease inhibitor mix, 10 mM sodium butyrate), as described. Following nuclei lysis, the extracts were precipitated with trichloroacetic acid, the pellets were resuspended in acetone supplemented with 0.1% HCl, and the isolated histones were sent for mass spectrometry analysis (IMSERC, Northwestern University).

#### 4tU-Seq and RNA half-life calculations

The 4tU-Seq and RNA half-life calculations were performed as described.^90^ Briefly, WT or *cct1-2* yeast were grown to log phase at 30°C, shifted to 37°C for 4 h, treated with either 4-thiouracil (4tU, Sigma Aldrich; 5 mM final) or DMSO for 5 min. Spike-in control cells, untreated *S. cerevisiae* or *K. lactis* cells treated with 4tU for 5 min, were added to the sample before RNA extraction. Following poly(A) RNA enrichment, fragmentation, and repair, the RNA was biotinylated with activated disulfide methane thiosulfonate (MTS) biotin (Biotium Biotin-XX MTSEA). Biotinylated RNA was collected using glycogen blocked MyOne Streptavidin C1 Dynabeads (Invitrogen). Following sequencing, the duplicate reads were removed and spike-in normalized nucleotide coverages were used to subtract background from the 4tU samples to obtain active transcription measurements. RNA half-life calculations were determined using the combined RNA-Seq and 4tU-Seq values.^65,67^ Briefly, the unitless constant of RNA half-life was estimated from the number of transcripts (RNA-Seq) divided by the production of new mRNA (4tU-Seq), with RNA-Seq and 4tU-Seq expressed as transcripts per million (TPMs).

#### Chromatin immunoprecipitation assay

Chromatin immunoprecipitations were performed as described.^60,91^ In general, WT or *cct1-2* yeast expressing the protein of interest as a TAP-tagged fusion were grown to log phase at 30°C, shifted to 37°C for 4 h, and collected for processing. Target proteins were immunoprecipitated using the TAP-tag as the epitope. Isolated DNA was analyzed by qPCR using primers select for each target DNA site, as marked. The relative DNA occupancies were determined using normalized PCR values relative to the control site *ACT1*.

#### Anchor-away and GFP-nanobody nuclear-depletion assays

To conditionally deplete yeast nuclei of the H2A.Z histone, we used a standard anchor away tactic.^58,92^ Briefly, the endogenous H2A.Z encoding gene (*HTZ1*) was engineered to express the protein as a carboxyl-terminal FRB fusion in a yeast background of *fpr1*Δ*tor1-1* (VDY1874).^93^ Rapamycin (1 μg/ml final) addition was used to trigger nuclear depletion or with ethanol addition as a control. To conditionally deplete yeast nuclei of TRiC, we exploited the strains expressing the Cct2 or Cct6 subunits as GFP fusions along with the inducible expression (GAL1p) of a GFP nanobody with nuclear export signal (NES). Transformants were grown in selective minimal media supplemented with raffinose (2%) to early log phase, galactose was added (2% final), and GFP signal was monitored by immunofluorescence.

#### TRiC protein purification

The TRiC complex was purified as previously described.^40^ In brief, High Five insect cells expressing His-tagged TRiC with GFP-Cct1 were collected and lysed by Dounce homogenization. The TRiC complex was captured by affinity purification using nickel resin. The TRiC complex was sequentially passaged over a heparin column, Mono Q ion exchange column, and Superose 6 size exclusion column. TRiC containing fractions were pooled, concentrated, and snap-frozen in liquid nitrogen.

#### Nuclear extract preparation

Functional nuclear extracts were prepared as described^94^, with slight modifications. Briefly, parental or *cct1-2* yeast grown at 37°C for 4 h were collected for spheroplasting and lysis by Dounce homogenization. The crude nuclei were lysed using 0.5 M ammonium sulfate and clarified using a SW55Ti rotor at 34100 rpm for 50 min at 4°C. Extracts were then clarified using a SW55Ti rotor at 12200 rpm for 15 min at 4 °C. The resulting pellet was resuspended and dialyzed with 3 changes of Buffer C (500 mL) supplemented with 75mM ammonium sulfate for a total of 4.5 h. Protein concentration of the nuclear extracts were determined using the BCA assay and stored in aliquots for in vitro transcription.

#### In vitro transcription assay

The *VAS1* (+100 to −50) DNA template for the in vitro transcription reactions was amplified from genomic DNA isolated from wild type yeast using an upstream primer with a biotin-labeled to allow immobilization on streptavidin metal beads. The in vitro transcription reactions contained immobilized template (100 ng of each template) and nuclear extracts (100 μg) prepared from either wild type (WT) or *cct1-2* yeast grown to log phase, shifted to 37°C for 4 h, and then collected. The cell pellets were used to prepare nuclear extracts as described.^94^ RNA transcripts were detected using standard primer extension reactions and radiolabeled primers designed to detect the upstream (aberrant) or downstream (coding) transcripts.^94,95^

## Acknowledgments

We are grateful to Janhavi Kolhe (UIUC), Lindsey Behrens (UIUC), and Anna Mankovich (UIUC) for their assistance and advice with the presented work.

## Author contributions

Z.G. designed the research, performed the research, analyzed the data; A.Y.T.P. performed the research; B.Z. analyzed the data; K.S. analyzed the data and secured funding; D.G. performed the research; J.F. analyzed the data and secured funding; B.C.F. designed the research, performed the research, analyzed the data, wrote the paper, secured funding.

## Inclusion and Diversity Statement

We support inclusive, diverse, and equitable conduct of research.

---

**Figure S1.**
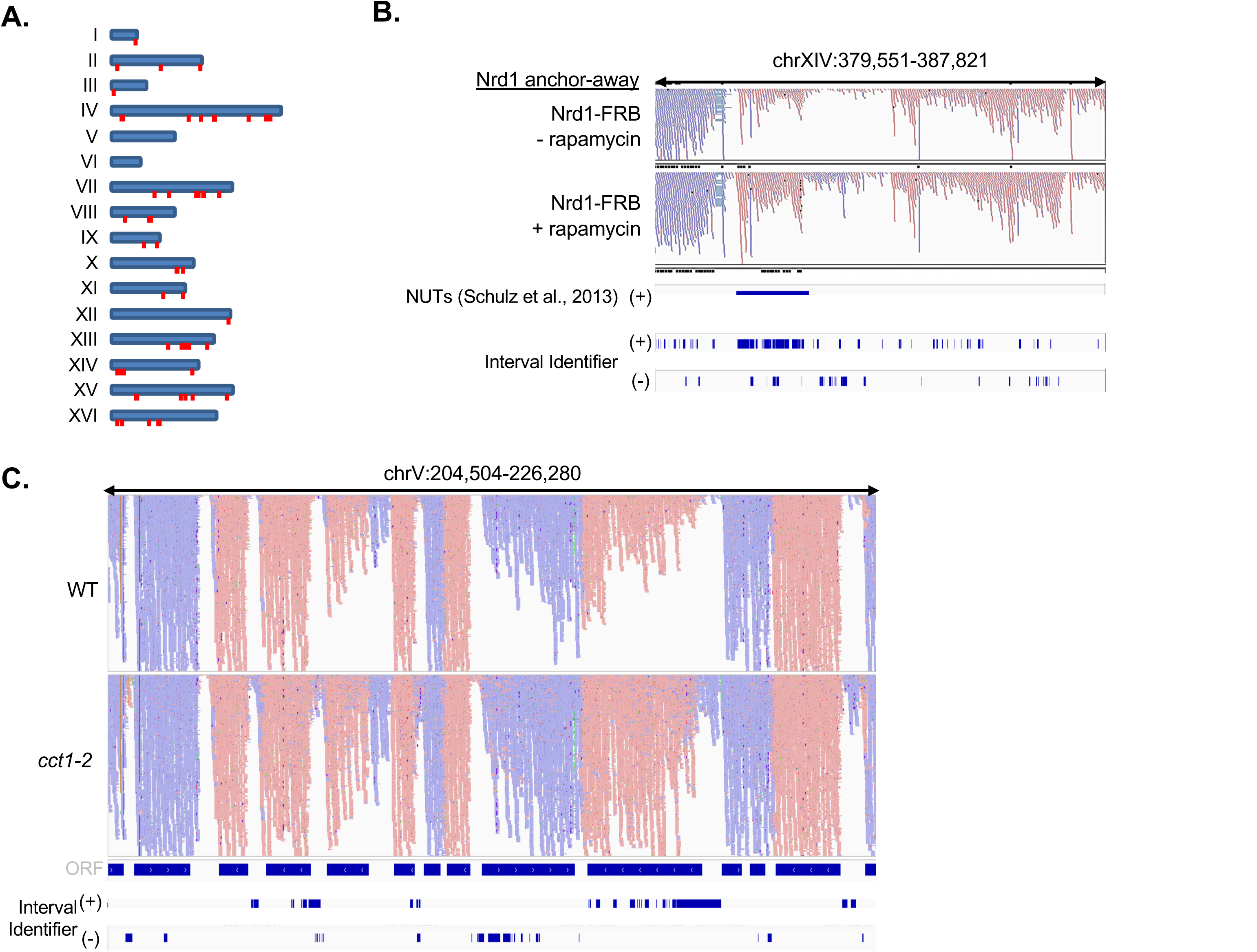
Genomic position of antisense loci and Interval Identifier reveals differences in aligned sequencing read counts. (**A**) The genomic position of gene loci predominantly transcribed in the antisense direction is shown. (**B**) The Interval Identifier algorithm effectively finds previously discovered cryptic RNAs. Shown is a comparison between an established NUT discovery and the results of Interval Identifier that shows the changes in read counts for both DNA strands, marked with (+) and (-). (**C**) Interval Identifier provides an in-depth analysis of read count variations between genomic data sets.

**Figure S2.**
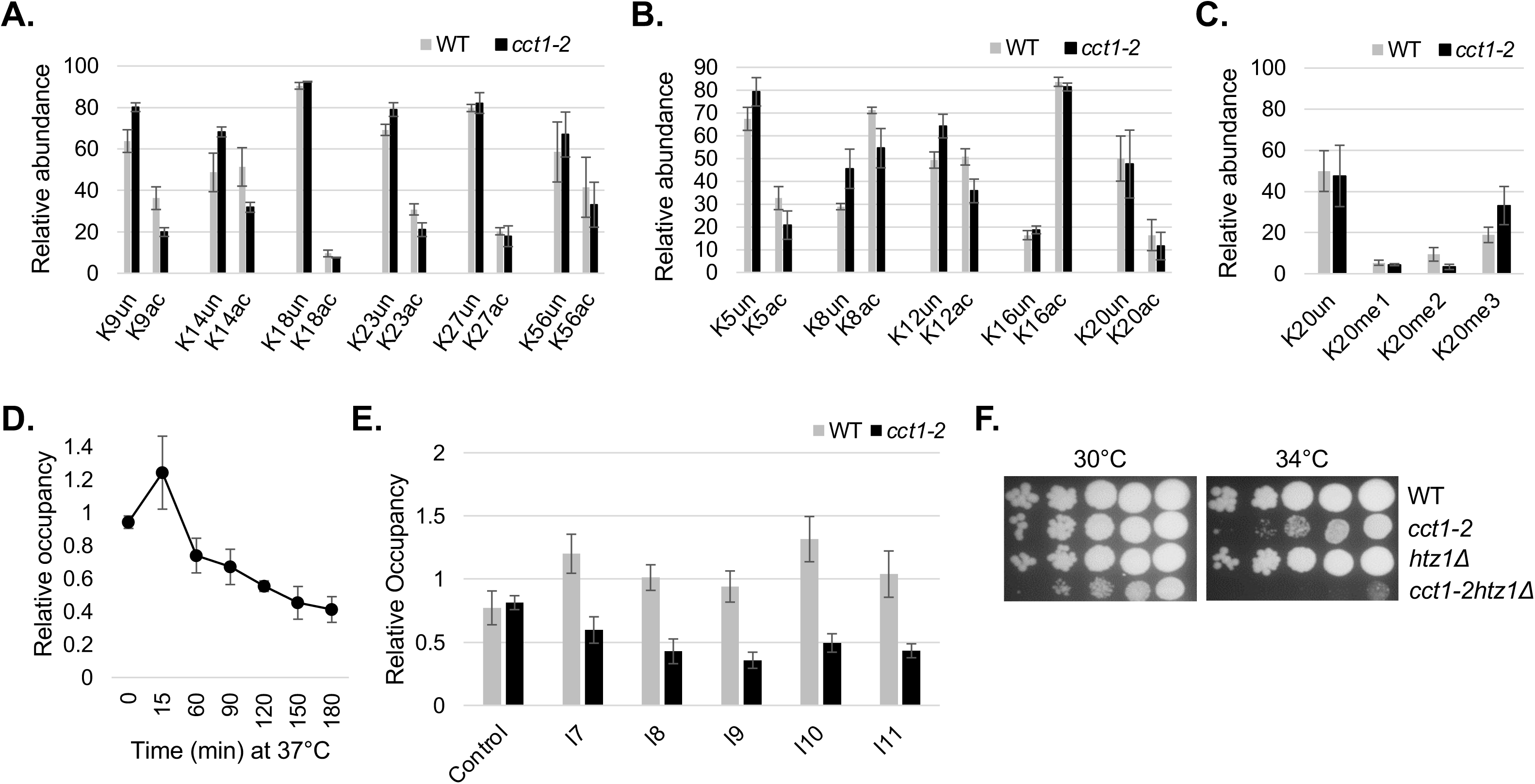
TRiC inactivation is not impactful on most histone posttranslational modifications yet TRiC is associated with H2A.Z. Mass spectrometry analysis of histone tail post-translation modifications was performed on histone proteins isolated from exponentially dividing wild type (WT) and *cct1-2* yeast exposed to 37°C for 4 h. The relative abundance of acetylation marks to the tail of histone 3 (**A**), histone 4 (**B**), and methylation of H4K20 (**C**) is shown. (**D**) The time-dependent, relative chromatin occupation of TAP-H2A.Z at intergenic region I8 was followed in *cct1-2* yeast at 30°C (0 min) and at the indicated times following a shift to 37°C, as marked. (**E**) The relative chromatin occupancy of TAP-H2A.Z was determined at control (ACT1) and I7-I11 intergenic regions *cct1-2* yeast after 4 h at 37°C. (F) TRiC and H2A.Z display a genetic interaction as determined using the spot test assay. Exponentially growing parental (WT), *cct1-2*, *htz1*1′, and *cct1-2htz1*1′ yeast were serially diluted, spotted onto media, and incubated at 30°C and 34°C, as indicated.

