## Supplemental Figures for "TRiC/CCT Chaperonin Governs RNA Polymerase II Activity in the Nucleus to Support RNA Homeostasis"

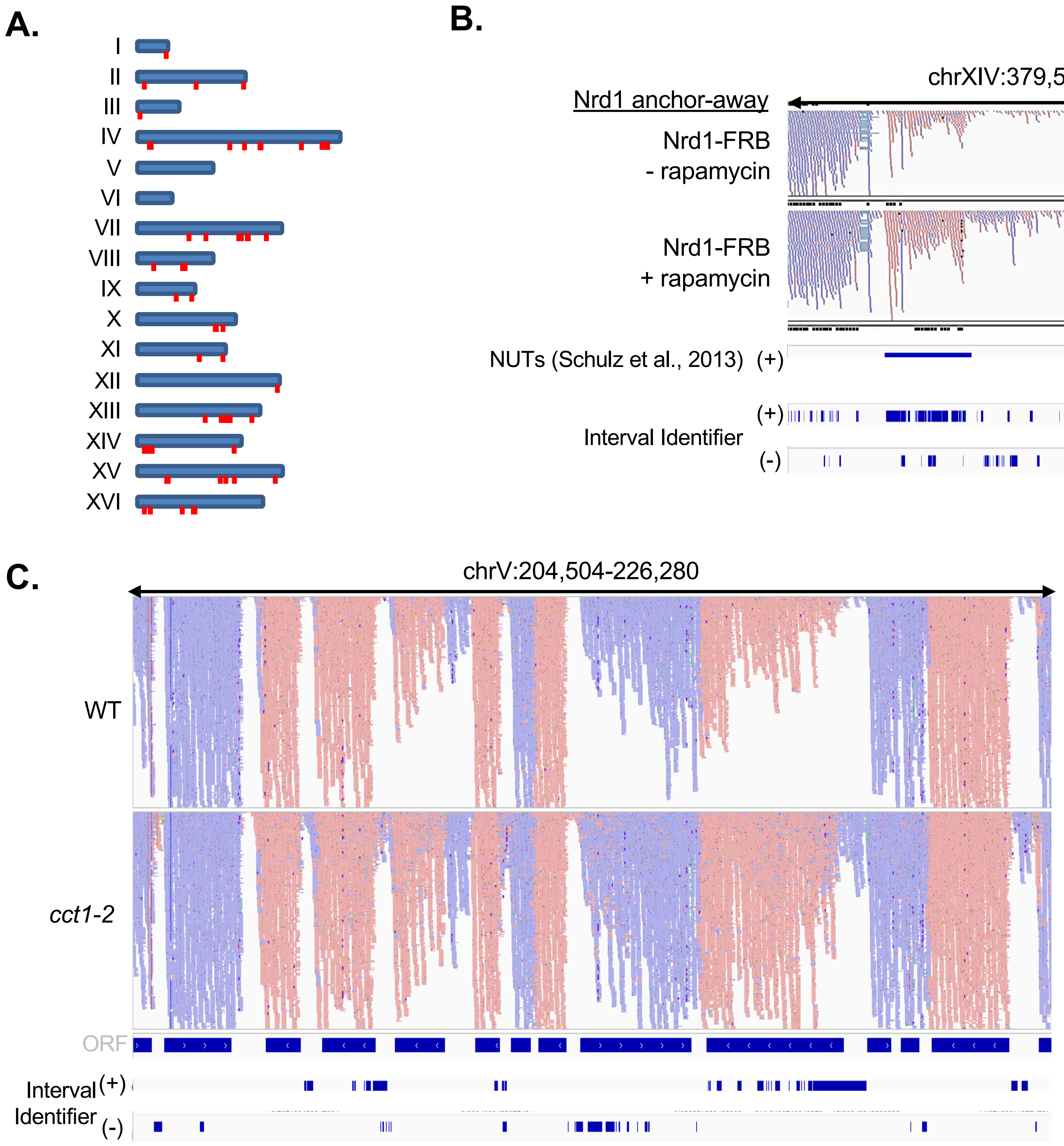


**Figure S1. Genomic position of antisense loci and Interval Identifier reveals differences in aligned sequencing read counts.** (**A**) The genomic position of gene loci predominantly transcribed in the antisense direction is shown. (**B**) The Interval Identifier algorithm effectively finds previously discovered cryptic RNAs. Shown is a comparison between an established NUT discovery and the results of Interval Identifier that shows the changes in read counts for both DNA strands, marked with (+) and (-). (**C**) Interval Identifier provides an in-depth analysis of read count variations between genomic data sets.

**Figure S2**

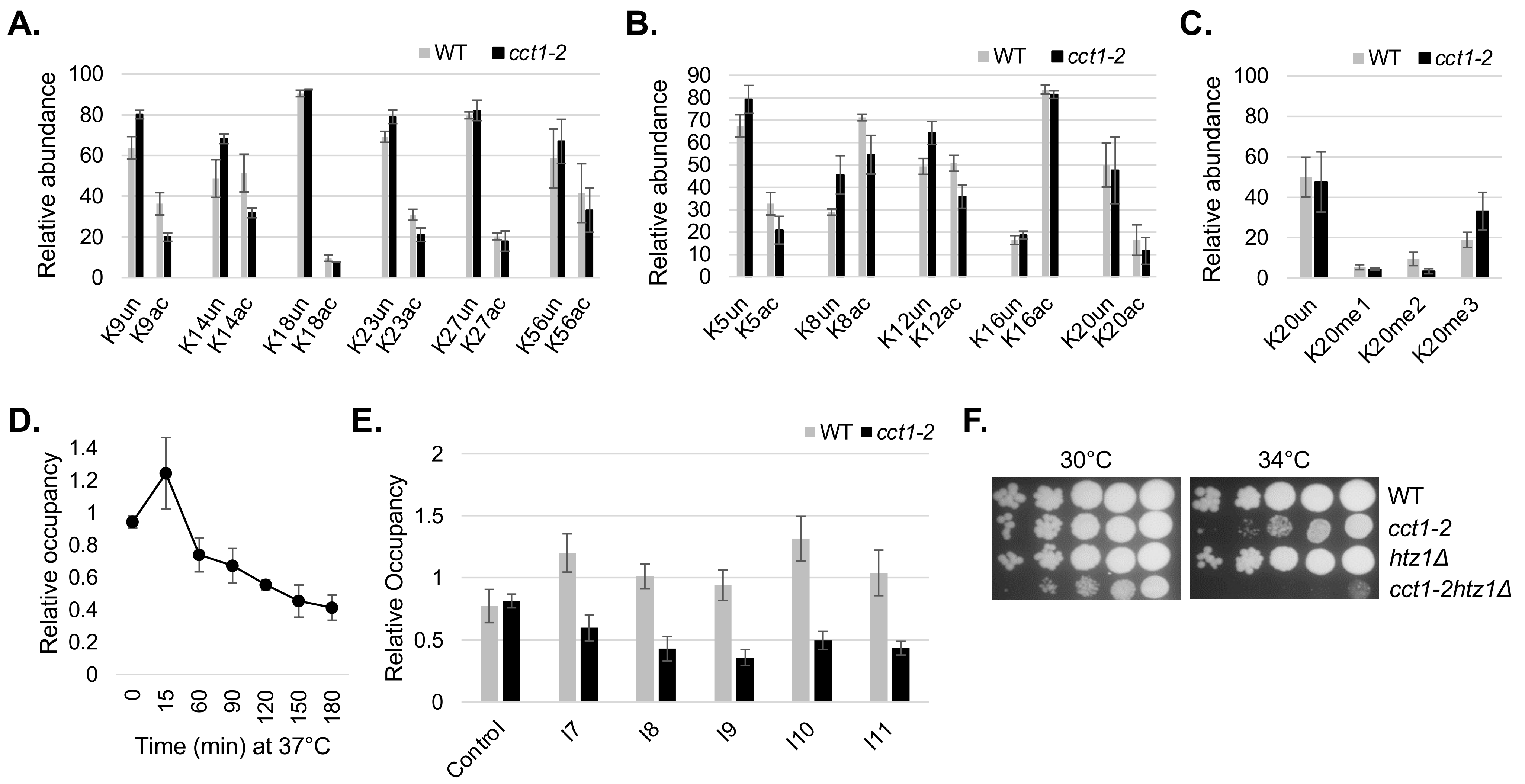


**Figure S2. TRiC inactivation is not impactful on most histone posttranslational modifications yet TRiC is associated with H2A.Z.** Mass spectrometry analysis of histone tail post-translation modifications was performed on histone proteins isolated from exponentially dividing wild type (WT) and *cct1-2* yeast exposed to 37°C for 4 h. The relative abundance of acetylation marks to the tail of histone 3 (**A**), histone 4 (**B**), and methylation of H4K20 (**C**) is shown. (**D**) The time-dependent, relative chromatin occupation of TAP-H2A.Z at intergenic region I8 was followed in *cct1-2* yeast at 30°C (0 min) and at the indicated times following a shift to 37°C, as marked. (**E**) The relative chromatin occupancy of TAP-H2A.Z was determined at control (ACT1) and I7-I11 intergenic regions *cct1-2* yeast after 4 h at 37°C. (F) TRiC and H2A.Z display a genetic interaction as determined using the spot test assay. Exponentially growing parental (WT), *cct1-2*, *htz1*Δ, and *cct1-2htz1*Δ yeast were serially diluted, spotted onto media, and incubated at 30°C and 34°C, as indicated.
